# Towards establishing internal validity for correlated gene expression measures in imaging genomics of functional networks: why distance corrections and external face validity alone fall short. Reply to “Distance is not everything in imaging genomics of functional networks: reply to a commentary on Correlated gene expression supports synchronous activity in brain networks”

**DOI:** 10.1101/2019.12.09.869412

**Authors:** Spiro P. Pantazatos, Mike Schmidt

**Affiliations:** Molecular Imaging and Neuropathology Division, New York State Psychiatric Institute, Department of Psychiatry, Columbia University Irvine Medical Center, New York, NY

## Abstract

The primary claim of the Richiardi et. al. 2015 Science article^1^ is that a measure of correlated gene expression, significant strength fraction (SSF), is related to resting state fMRI (rsfMRI) networks. However, there is still debate about this claim and whether spatial proximity, in the form of contiguous clusters, accounts entirely, or only partially, for SSF^2,3^. Here, thirteen distributed networks were simulated by combining 34 contiguous clusters randomly placed throughout cortex, with resulting edge distance distributions similar to rsfMRI networks. Cluster size was modulated (6-15mm radius) to test its influence on SSF false positive rate (SSF-FPR) among the simulated ‘noise’ networks. The contribution of rsfMRI networks on SSF-FPR was examined by comparing simulations using: 1) all cortical samples 2) all samples with non-rsfMRI cluster centers and 3) only non-rsfMRI samples. Results show that SSF-FPR is influenced only by cluster size (r>0.9, p<0.001), not by rsfMRI samples. Simulations using 14mm radius clusters most resembled rsfMRI networks. When thresholding at p<10^-4^, the SSF-FPR was 0.47. Genes that maximize SF have high *global* spatial autocorrelation. In conclusion, SSF is unrelated to rsfMRI networks. The main conclusion of Richiardi et. al. 2015 is based on a finding that is ∼50% likely to be a false positive, not less than 0.01% as originally reported in the article^1^. We discuss why distance corrections alone and external face validity are insufficient to establish a trustworthy relationship between correlated gene expression measures and rsfMRI networks, and propose more rigorous approaches to preclude common pitfalls in related studies.

## Introduction

There is still active debate about whether brain regions comprising resting state fMRI networks exhibit uniquely high correlated gene expression or whether this effect is attributed entirely to spatial proximity in the form of contiguous clusters^1–3^. The main claim and conclusion of the Richiardi et. al. 2015 Science article^1^ is that high correlated brain gene expression (i.e. significant strength fraction (SSF)) can be at least partially explained by and is *related* to resting state fMRI (rsfMRI) networks. Our 2017 commentary^2^ found that randomly spaced clusters also generate SSFs much higher than expected by chance, indicating that SSF is *not* specific to rsfMRI networks. On the basis of this and other evidence presented in our 2017 commentary^2^, we claimed that SSF is attributable entirely to spatial proximity, and that it is unrelated to rsfMRI networks. In their reply to our commentary, Richiardi et. al. 2017^3^ did not rebut this claim, other than to (rightly) point out that edges in these random clusters (median < 24 mm) were shorter than rsfMRI networks (median < 50 mm). That the SFs tend to be higher overall in these simulated networks (see Figure 1D in our 2017 commentary), which also tend to have shorter edges, is consistent with our argument that distance drives SSF. However, it is unclear, *a priori,* why lower average within-network (W) distances could account for the non-specificity of SSF. More rigorous and comprehensive network ‘noise’ simulations are needed to determine the impact of spatial proximity (i.e. contiguous cluster size) vs. rsfMRI networks on SSF and whether SSF can be attributed entirely, or only partially, to spatial proximity.

**Figure 1.**
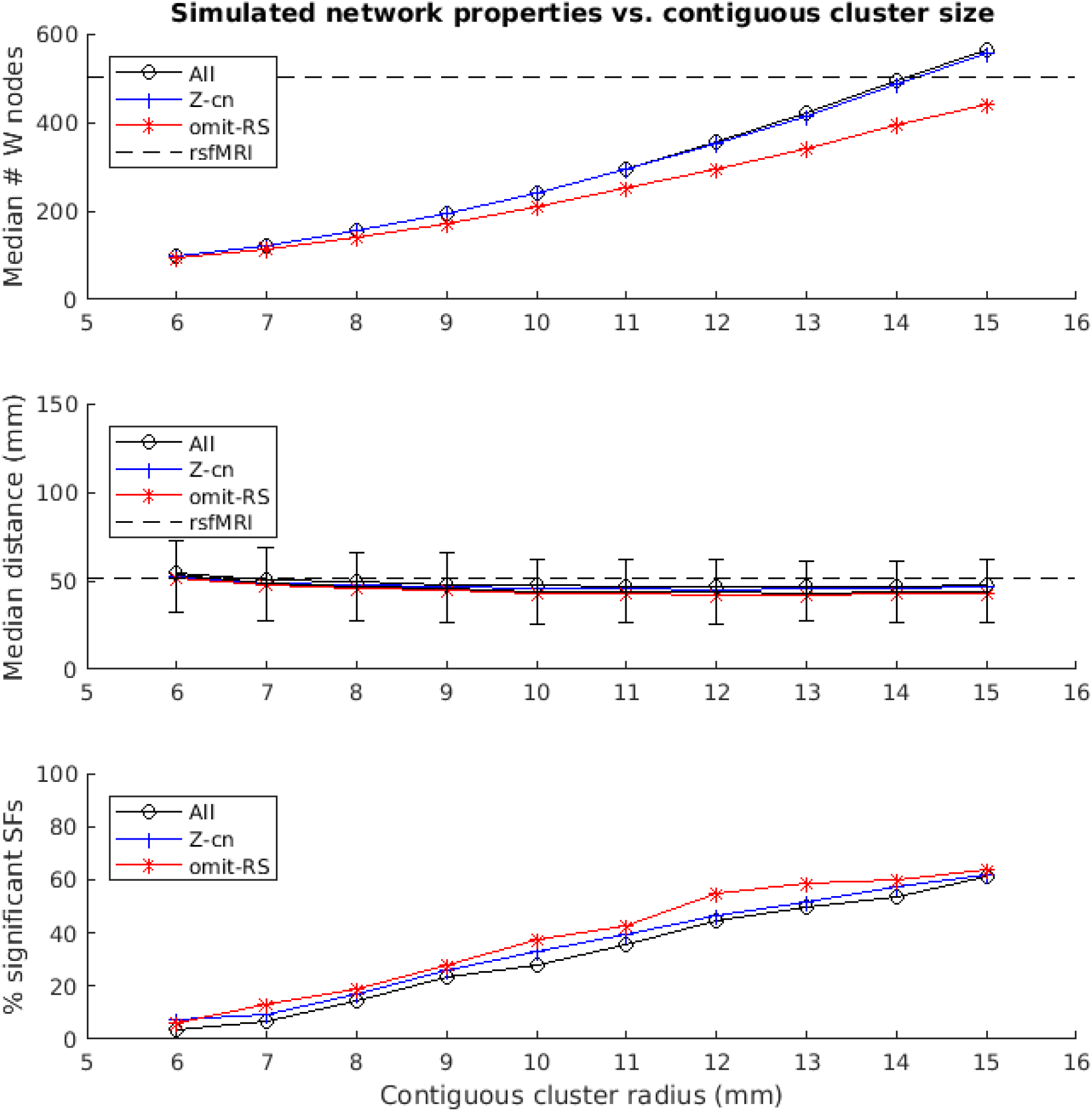
Plots of simulated network properties vs. contiguous cluster size. **Top panel**) Median network sample size (W). Dashed line depicts the value for real rsfMRI networks (W=501). **Middle panel**) Median network edge distances (90% of the median distances across the 1,000 random networks fell within the depicted error bars). Dashed line depicts the value for real rsfMRI networks (<50 mm). **Bottom panel**) Contiguous cluster size, not rsfMRI samples, predicts percentage of SSF (SSF-FPR x 100) among the simulated ‘noise’ networks (All: r=0.98; Z-cn: r=0.99; omit-RS: r=0.99, p-values<10E-6). **All** = experiment in which all 1,777 AHBA cortical samples were used; **Z-cn** = same as “All”, except cluster centers were forced to line outside of rsfMRI network areas; **omit-RS** = only non-rsfMRI samples (1,276) were used to simulate both network and non-network regions.

Here, we generated randomly distributed networks so that they more accurately simulate rsfMRI networks. The parameters of the simulations were varied in order to examine the impact of contiguous cluster size on frequency of SSF among the simulated networks (i.e. false positive rate of SSF, or SSF-FPR). When using all samples for each network simulation, contiguous cluster centers have <20% chance of being a rsfMRI sample (note this is a rough estimate since the probability is 28% (501/1777) when the first cluster is formed and then drops slightly with subsequent clusters because they are not allowed to overlap with any previously formed clusters, see methods). Therefore, an additional variation of simulations was run which enforced non-rsfMRI cluster centers. To further the reduce the chances that simulated clusters overlapped with and included *any* rsfMRI samples, a third variation removed them entirely from the simulations. *If rsfMRI networks are related with and contribute at all towards SSF, then the SSF-FPR should decrease with the latter two experiment variations*.

In their reply to our commentary, Richiardi et. al. claimed their results hold after linear distance correction (based on regression). We plotted tissue-tissue correlations vs. distance to show why this approach is insufficient to remove the effects of spatial proximity. Finally, we propose that the feature selection routine applied in the original 2015 article selects the genes that exhibit the highest *global* spatial autocorrelation, consistent with our argument that spatial proximity alone drives SSF. We tested this by examining and comparing measures of spatial autocorrelation for the 136 consensus features identified in the original 2015 article vs. all other genes.

## Methods

A series of networks were simulated for varying sizes of contiguous clusters (i.e. 6-15 mm radius spheres) placed randomly throughout cortex. Cluster centers were comprised of cortical samples chosen at random from among the 1777 included in the original Richiardi et. al. Science article. For these networks, 34 randomly placed, contiguous and non-overlapping clusters were first generated, then combined to form 9 distributed ‘networks’ comprised of 3 clusters, 3 networks comprised of 2 clusters, and one network comprised of a single cluster.

For each of the 10 sizes, 1,000 distributed networks were simulated, using 1,000 shuffles (permutations) for SF significance testing, yielding 10,000,000 SF calculations (iterations) per experiment variation. SFs were considered significant at p < 1/(# total shuffles), or 0.001 in this case. The simulations were repeated for each the 3 experiment variations: 1) “All” - using all cortical samples, 2) “Z-cn” - using all cortical samples but enforcing non-rsfMRI cluster centers and 3) “omit-RS”- using *only* non-rsfMRI samples. The total # of available samples for the above 3 experiment variations were 1777, 1777 and 1276, respectively. The same procedures, including within tissue correction etc., were applied as in the original 2015 article and in our 2017 commentary when calculating SF.

Moran’s I, a measure of spatial autocorrelation, was calculated for each of 16,906 genes’ expression levels. Moran’s I approaches +1 when the gene expression levels cluster perfectly over space (high spatial autocorrelation), and approaches −1 when the tested variable is evenly distributed (high dispersion). A measure of 0 indicates no relationship between gene expression levels are randomly dispersed in space. Moran’s I was assessed with pysal v2.0 libraries^4^, using continuously diminishing weights over distance from each sample location up to 16mm, beyond which all further weights were set to 0. Neither changing the 16mm threshold to 64mm nor using binary rather than continuous weighting substantively changed the results. The updated MATLAB code (network simulations and resulting plots and figures) and a Jupyter notebook and Python code (gene expression spatial autocorrelation analyses and resulting plots) are available at https://github.com/spiropan/ABA_functional_networks.

## Results

A contiguous cluster size of 14 mm yielded within network sample size (W) closest to the rsfMRI sample size of 501 (Figure 1, top panel). Varying contiguous cluster size did not significantly impact median within network (W) edge distances of the random networks, which all approximated the median edge distance of real rsfMRI networks (dashed line in Figure 1, middle panel). Contiguous cluster size almost perfectly predicted SSF-FPR (r values>0.98, p<10E-6, Figure 1, bottom panel). The SSF-FPR rose from <.05 at 6 mm radius to >0.6 at 15 mm (in other words, about 50x to 600x the theoretical FPR at alpha level p=0.001). Forcing cluster centers to lie outside rsfMRI areas or even removing rsfMRI samples all together did not impact the results In these simulations, the latter actually *increased* the SSF-FPR at most cluster sizes (Figure 1, bottom panel). An additional simulation at cluster size 14mm, this time using 10,000 (vs. 1,000) shuffles for SF significant testing, yielded SSF-FPR=0.47 (i.e. 4,700x higher than the theoretical false positive rate p<10^-4^ reported in the original 2015 Science article). Note our approach simulated 13 networks that roughly mimic the distribution of median distances of the rsfMRI networks shown in Figure 1 of the 2017 reply from Richiardi et. al^3^. Figure 2A here recapitulates that figure in boxplot form and also shows an example set of simulated networks with cluster size=14mm in the current study underneath.

**Figure 2.**
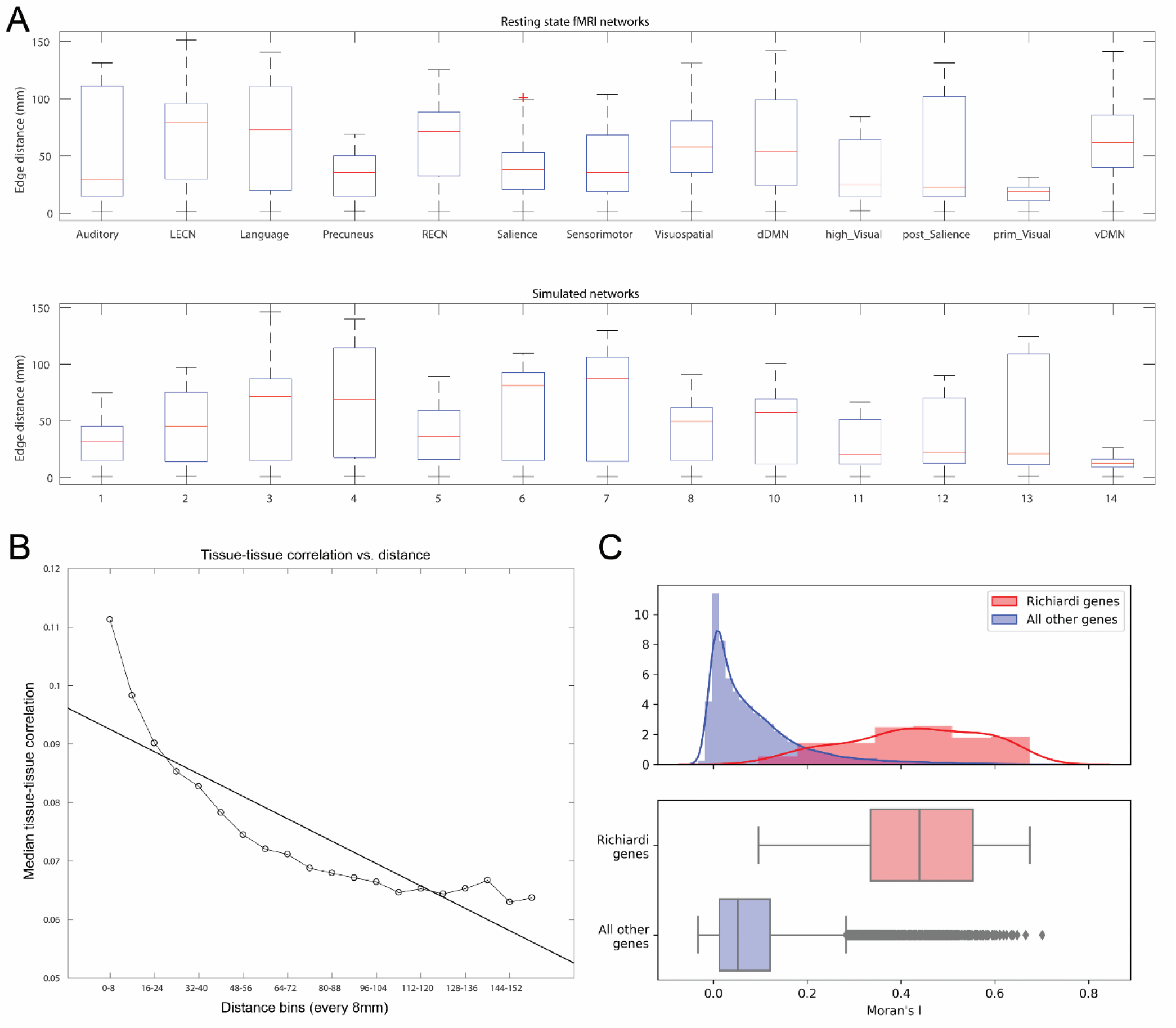
**A.** Boxplot of edge distances (y-axis) for resting state fMRI networks as defined in Richiardi et. al. 2015 (top panel), and an example set of simulated networks in the current study (bottom panel). **B.** A plot of distance (8 mm bins, X-axis) vs. median brain gene expression tissuetissue correlation (y-axis). The plot used all edges used to calculate SF after applying within-tissue correction. **C.** Consensus features that maximize high strength fraction (identified in the original 2015 article) are also more likely to cluster together in space and exhibit high global spatial autocorrelation. Top panel: Histograms and kernel density plots of Moran’s I values for each of 16,906 genes, 136 high SF genes in red, all others in gray. Bottom panel: The y-axis is relative to each sample, with different n, so heights are not comparable, but position along the x-axis is. Bottom panel: A box plot of the same Moran’s I values.

A control analysis similar to the “All” condition tested whether total # W samples (not contiguous cluster size) predicted SSF. To reduce computation time, only 200 networks were simulated using 200 shuffles for SF significance testing. The contiguous cluster size was held constant at 6mm, while scaling factors from [1, 1.5,2.·.5] modulated the number of contiguous clusters comprising the networks. While this varied the median # W nodes from <100 to 450 among the simulated networks (median edge distances <57 mm), the SSF-FPR *decreased* slightly from 0.12 to 0.085 (r=-0.2, p=0.6, data not shown), confirming that contiguous cluster size, and not total # W nodes, predicts SSF.

We plotted median tissue-tissue correlations (after applying within-tissue correction as in the original 2015 article) vs. distance to confirm a non-linear relationship (Figure 2B). Next, we tested whether the 136 consensus features identified in the original 2015 article exhibit high global spatial autocorrelation. The relationship between the magnitude of each gene’s expression and its spatial clustering was assessed using Moran’s I, a measure of spatial autocorrelation. Moran’s I was significantly higher in the 136 genes Richiardi deemed significant (mean I = 0.430, sd 0.142) than in remaining genes (mean I = 0.084, sd 0.100) (p < 10^-9^ for each of the 136 genes). See Figure 2C.

## Discussion

The main finding and claim of the original 2015 article, reflected in the title, is “that functional brain networks defined with resting-state functional magnetic resonance imaging can be recapitulated by using measures of correlated gene expression in a post mortem brain tissue data set.” The main text states “the spatial organization of functional networks corresponded to regions that have more highly correlated gene expression than expected by chance (*P* < 10^-4^)” and that “this finding cannot emerge from spatial proximity or gross tissue similarity”. This is the most critical, specific claim under contention.

Our network simulations indicate that the spatial organization of rsfMRI networks **do not** correspond to regions that have more highly correlated gene expression than expected by chance. The probability of observing highly correlated gene expression for rsfMRI networks by chance is about 50%, not less than 0.01% as reported in the original article. Furthermore, contiguous cluster size, and not resting state fMRI networks, accounts entirely for SSF. The SSF-FPR was not affected by enforcing non-rsfMRI cluster centers or removing rsfMRI samples entirely from the simulations. Taken together, these results indicate that the evidence presented in the original 2015 article is insufficient to establish a relationship between correlated brain gene expression and rsfMRI networks. In other words, SSF is not an internally valid measure.

### Why distance corrections alone are not enough to establish links between correlated brain expression and rsfMRI networks

In their reply to our 2017 commentary, Richiardi et. al. rightly point out that Euclidean distance correction will ‘wrongly assign “nearness” to two “neurally distant” regions on the crowns of adjacent gyri’ (see figure in their reply). This is one reason why correcting for Euclidean distance alone is not adequate and why other types of control experiments are required to validate measures of correlated gene expression such as SSF. In their reply, Richiardi et. al. also argue that their original results hold after distance corrections using linear regression and distance-aware permutation. Below we discuss why these approaches alone are not enough to validate the SSF measure.

Our commentary discusses why Euclidean distance (linear regression) is an inadequate method for proximity correction due to strong model assumptions (i.e. the relationship between distance and tissue correlation is not linear, as evidenced by a plot of median tissue-correlations vs. distance, Figure 1). The authors suggested an intrinsic contradiction in our recommendation to avoid distance correction using linear regression and then also showing a linear correlation in our original 2017 commentary Figure 1B. There was no contradiction here, because a best fit line was added only to show the first order approximate fit between distance and correlated gene expression, not to suggest linear regression be used for distance correction. A primary purpose of Figure 1B was to show that the within-network (Wi) edges (dark grey) are substantially shorter than the out of network edges (T-Wi, light gray). Our critique was also founded on Figure 1A (removing within tissue samples inadequately corrects for spatial proximity), Figure 1C (SF falls monotonically as progressively longer edges are removed) and Figure 2D (SSF is not specific to resting state fMRI). In hindsight, we could have tried to overlay the curve (shown in Figure 1 here) to our 2017 commentary Figure 1B, but it would have been less straightforward to attach a p-value and the model fit might not have been readily apparent.

The distance-aware permutation testing appears to be a more valid approach than the original (non-distance) aware permutation testing. However, the authors’ results (higher p-values with more conservative distance aware corrections) illustrate and support the fact that SF is driven by distance, but in a different way that parallels our 2017 commentary Figure 1C. Whereas our 2017 commentary Figure 1C showed that ‘real’ SF ***decreases*** as ***short*** edges are removed from the rsfMRI networks, the author’s results show that the null distribution SFs ***increase*** as ***long*** edges are removed from the ‘null’ networks. Both scenarios will create higher p-values in SF significance testing.

Critically, the fact the authors’ new analyses survive distance corrections and distance aware permutations, and at higher p-values than <10^-4^ as originally reported in the 2015 article, does not validate the SSF measure, since it is unrelated to rsfMRI networks to begin with. In other words, these same distance corrections and distance aware permutations would show similar results (i.e. higher SF p-values but still less than 0.01 or 0.05) when applied to any of the <50% of randomly spaced networks that started out with SF p-values less than 10^-4^.

### Why face validity in independent datasets is not enough to establish a link between SSF and rsfMRI networks

In response to our 2017 commentary, Richiardi et. al. stressed their replications (more appropriately called face validity) in independent datasets and note that we did not generate gene lists for any of our random cluster analyses and examine them in other independent datasets. There are several problems with this line of reasoning and argumentation. First, the authors’ response appears to be “moving the goalpost”^1^. The authors did not rebut our 2017 specific claim that SSF is both unrelated and not specific to rsfMRI networks, other than to rightly note that the median distances of our simulated ‘noise’ networks were half as long as rsfMRI networks. To rebut our claim, the authors would have needed to show that simulated networks with longer median distances fail to generate inflated SSF-FPR, but they did not.

Secondly, we did not claim that the set of 136 genes identified in the study were not important to functional connectivity. However, *at best,* the additional findings that the authors mention in their reply (i.e. in mouse connectivity and resting state fMRI connectivity using their identified set of 136 genes), suggest *face validity* for, but not evidence that *validates,* their primary

It is plausible that, even if SSF is non-specific and completely unrelated to rsfMRI networks when using all genes, the 136 consensus features identified in the original 2015 article are unique to rsfMRI networks and uniquely important to functional connectivity. To demonstrate this would require additional control experiments and comparisons. One possibility is that the consensus feature selection routine used in the original 2015 article converges on genes that tend to be more highly expressed in the largest contiguous clusters of any network. In the case of rsfMRI, these include the posterior cingulate and ventromedial PFC of the dDMN, which are major hub regions and highly connected to the rest of the brain, which could help explain the evidence for supporting connectivity in independent datasets. Critically, the independent replication (more appropriately called external face validity) noted by Richiardi et. al. in their reply, (https://arxiv.org/abs/1706.06088), used *all* genes as a baseline group when examining the 136 consensus features identified in the original 2015 article. A more rigorous baseline group would have been all genes with properties similar to the 136 genes (i.e. mean expression in the brain, variability and global spatial autocorrelation) in order the confirm that these generic properties alone do not account for observed relationships with connectivity.

It is also possible that the identified consensus features *are not* unique to rsfMRI networks. This interpretation is supported by our results that found the 2015 consensus features have high global spatial autocorrelation throughout cortex (Figure 2C). It is also possible that the 136 consensus features *are* relatively unique to rsfMRI networks, but that consensus features identified in random networks are *also* important to functional connectivity and demonstrate face validity. Further work is required to determine which scenario is true. Any of the above scenarios, however, is independent of the primary claim of the original 2015 article.

### Suggestions for future work

When dealing with high-dimensional data, it is difficult to isolate relevant variables in light of the fact that any set of features can be identified when maximizing some variable or measure, even with ‘noise’ data. It is also important to first ensure that the measure and its interpretation is *internally valid,* before seeking external validations. Here, network ‘noise’ simulations were used to test the internal validity of the SSF measure and its interpretation that it is specifically related to rsfMRI networks. Future studies of imaging genomics and functional networks that use measures of correlated gene expression could combine non-linear or non-parametric distance correction, distance-aware permutation as well as split-half bootstrap resampling to rigorously test the validity of the measures and their interpretation and also the consistency of the features (genes) that optimize the measures.

## Conflict of Interest Statement

The authors declare that the research was conducted in the absence of any commercial or financial relationships that could be construed as a potential conflict of interest.

## Acknowledgments

This work was supported by a K01MH108721 (SPP). We would like to thank Paul Pavlidis for helpful comments and suggestions.

## Author Contributions

S.P.P conducted network simulation analyses and drafted the manuscript, M.S conducted spatial autocorrelation analyses and drafted relevant portions of the text.

## Footnotes

1 “Moving the goalposts is an informal [logical] fallacy in which evidence presented in response to a specific claim is dismissed and some other (often greater) evidence is demanded.” (https://en.wikipedia.org/wiki/Moving_the_goalposts) claim, which is that correlated gene expression is higher in and uniquely related to resting state fMRI networks.

## References

1. Richiardi, J. et al. BRAIN NETWORKS. Correlated gene expression supports synchronous activity in brain networks. Science 348, 1241–1244 (2015).

2. Pantazatos, S. P. & Li, X. Commentary: BRAIN NETWORKS. Correlated Gene Expression Supports Synchronous Activity in Brain Networks.Science348, 1241-4. Frontiers in neuroscience 11, 412–412 (2017).

3. Richiardi, J., Altmann, A. & Greicius, M. Distance is not everything in imaging genomics of functional networks: reply to a commentary on Correlated gene expression supports synchronous activity in brain networks. bioRxiv 132746 (2017) doi:10.1101/132746.

4. Rey, S. J. & Anselin, L. PySAL: A Python Library of Spatial Analytical Methods. in Handbook of Applied Spatial Analysis: Software Tools, Methods and Applications (eds. Fischer, M. M. & Getis, A.) 175–193 (Springer, 2010). doi: 10.1007/978-3-642-03647-7_11.

